# A method to exploit the structure of genetic ancestry space to enhance case-control studies

**DOI:** 10.1101/043166

**Authors:** Corneliu A. Bodea, Benjamin M. Neale, Stephan Ripke, The International IBD Genetics Consortium, Mark J. Daly, Bernie Devlin, Kathryn Roeder

## Abstract

One goal of human genetics is to understand the genetic basis of disease, a challenge for diseases of complex inheritance because risk alleles are few relative to the vast set of benign variants. Risk variants are often sought by association studies in which allele frequencies in cases are contrasted with those from population-based samples used as controls. In an ideal world we would know population-level allele frequencies, releasing researchers to focus on case subjects. We argue this ideal is possible, at least theoretically, and we outline a path to achieving it in reality. If such a resource were to exist, it would yield ample savings and would facilitate the effective use of data repositories by removing administrative and technical barriers. We call this concept the Universal Control Repository Network (UNICORN), a means to perform association analyses without necessitating direct access to individual-level control data. Our approach to UNICORN uses existing genetic resources and various statistical tools to analyze these data, including hierarchical clustering with spectral analysis of ancestry; and empirical Bayesian analysis along with Gaussian spatial processes to estimate ancestry-specific allele frequencies. We demonstrate our approach using tens of thousands of controls from studies of Crohn’s disease, showing how it controls false positives, provides power similar to that achieved when all control data are directly accessible, and enhances power when control data are limiting or even imperfectly matched ancestrally. These results highlight how UNICORN can enable reliable, powerful and convenient genetic association analyses without access to the individual level data.

## 1 Introduction

To detect genetic variants affecting risk for complex disease, the ideal association study would contrast a large number of affected subjects to an even larger set of population-based samples used as controls. Ideally these controls would be so numerous and so well-matched to cases, ancestrally, that the power to detect risk variants would be limited solely by the size of the case sample. This article outlines an approach to turn this ideal into reality.

The challenges in accruing a large control sample are numerous. It requires a substantial portion of the research budget; while data repositories, such as dbGaP [1, 2], contain genetic data from tens of thousands of potential control samples, using these data requires considerable and independent effort from each research team; and issues such as population structure and genotyping platform require additional work before an adequately-controlled association test can be performed. Family-based studies obviate concerns about ancestry [3, 4], but they have other drawbacks [5, 6, 7, 8, 9].

Instead we show here that it is theoretically possible to build a web resource that enables research teams to focus on maximizing the value of their case sample, by providing control allele frequency information that is optimally matched to the available cases. Additionally, information can be exchanged via a web server similar to the existing Exome Aggregation server (exac.broadinstitute.org), without revealing individual genetic information. We call such a resource the Universal Control Repository Network (UNICORN), because it provides matched control data for a variety of ancestries. In our vision, and to ensure the confidentiality of both cases and controls, no case genotype information is passed to UNICORN, nor will the controls data processed to produce UNICORN be accessible to this resource.

Our approach to building UNICORN employs the spectral graph approach [10], which has similarities to principal component analysis [11, 12], to obtain a hierarchical representation of ancestry, where individuals are clustered into increasingly finer ancestry spaces. Using a Bayesian model we infer allele frequencies over all such clusters, always borrowing strength across the entire hierarchy to maximize power. We then perform a second layer of inference within clusters to model spatial variation. This step picks up fine-grained ancestry structure that the hierarchical clustering did not by assuming that deviations from a cluster-wide average follow a Gaussian process with a covariance structure that is inferred from the ancestry space. This model is appropriate because it is flexible enough to accommodate smooth allele frequency fluctuations with varying degrees of spatial correlation.

Our results on both simulated data and imputation-based genotype level data from seven studies of Crohn's disease show that UNICORN has the potential to greatly improve power in genetic association tests. First we show that UNICORN not only controls false positives, but it makes efficient use of the control data, providing power similar to a setting in which all control data are directly accessible to the researcher. We then show that UNICORN can improve power relative to a carefully matched case-control study simply by using all available control information, even though the additional controls are not perfectly matched to cases.

## 2 Subjects and Methods

### 2.1 Overview of UNICORN

The steps involved in building our version of UNICORN (henceforth simply UNICORN) and performing an association study are now outlined (Figure 1). Existing publicly available collections of control data determine a common genetic ancestry space onto which cases and controls can be projected independently. GemTools [10, 13, 14] constructs ancestry spaces and performs such projections. The projected controls are then used to estimate the control minor allele frequency distribution (MAFD) over the ancestry space. For efficiency of computations, the MAFD would be precomputed and stored for application whenever users request control information. To query the repository, researchers project their cases onto the public control ancestry space and submit the locations to UNICORN. Based on the pre-computed surface, the system will infer allele frequencies as well as the degree of uncertainty associated with the estimates at all relevant locations and return the results to the users, who can then proceed with an association test, such as the one we describe in the sequel.

**Figure 1:**
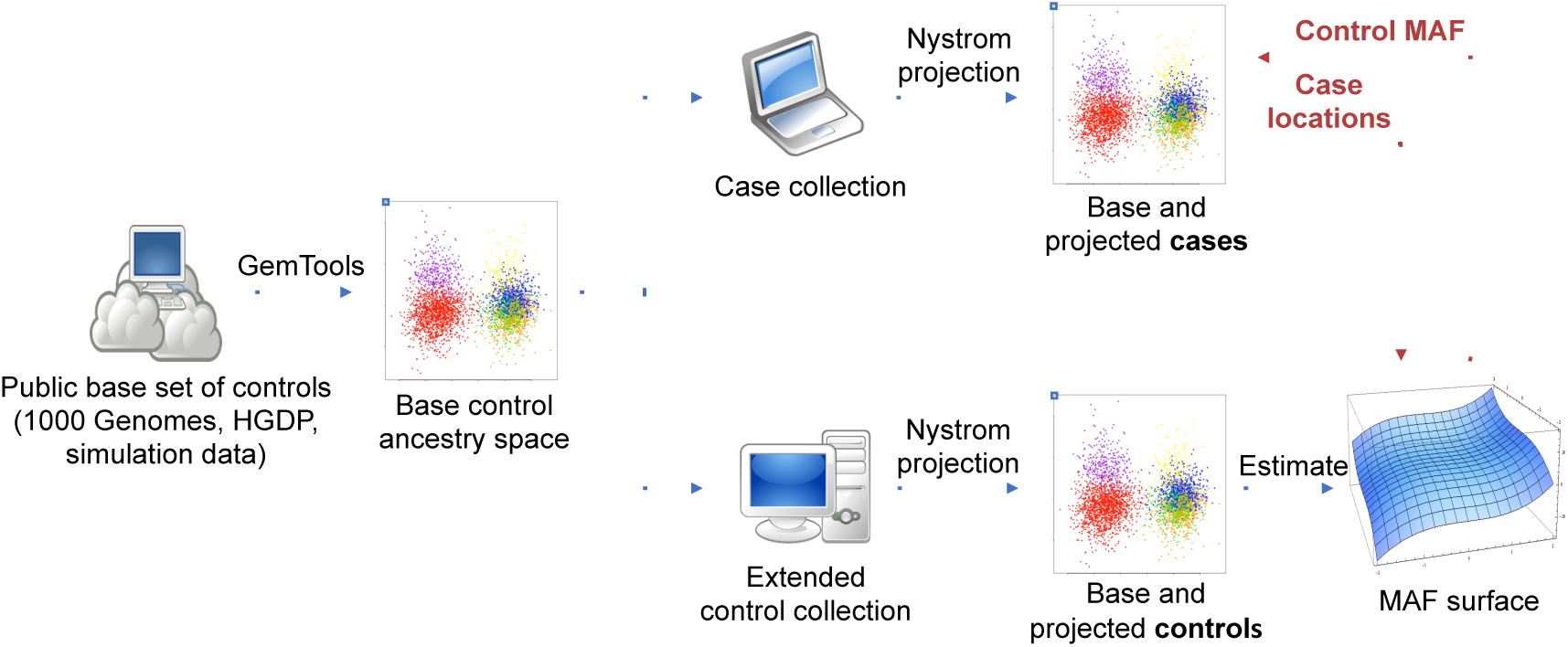
Overview of the UNICORN model. The UNICORN pipeline starts with a public base set of controls and constructs the corresponding base control ancestry space. All subsequent cases and controls can be projected independently via GemTools onto this space. This approach ensures that, having only knowledge of the base set, new individuals can be compared to existing ancestries. An extended set of controls is then projected onto the base control ancestry space, which is used to estimate the minor allele frequency distribution (MAFD) over the ancestry space. To query the repository, researchers project their cases onto the base control ancestry space and submit the resulting coordinates to the UNICORN server. Users then receive control allele frequencies as well as the degree of uncertainty associated with these estimates for all relevant locations, based on the pre-computed MAFD. Users can then proceed with an association test. Users only need to submit ancestry coordinates and the system only returns frequency inferences for the corresponding locations (red arrows). No other information is exchanged.

To estimate the MAFD, UNICORN employs a combination of empirical Bayesian analysis across a hierarchical clustering of the controls and, for localized ancestry regions, a Gaussian process model of the minor allele frequency (Figure 2). To visualize the algorithm in action we utilize the the Europeans in the Population Reference Sample (POPRES) [15] (dbGaP accession number phs000145.v4.p2), which yields an ancestry map that approximates the geographic map of Europe [16, 17]. Two SNPs in LCT (lactase persistence) and OCA2 (hair, skin and eye color) provide examples of UNICORN's MAFD for SNPs under selection, and provide an illustration of clines in allele frequency across Europe. Intensity of color displays allele frequency estimates that vary smoothly across the map (Figure 3).

**Figure 2:**
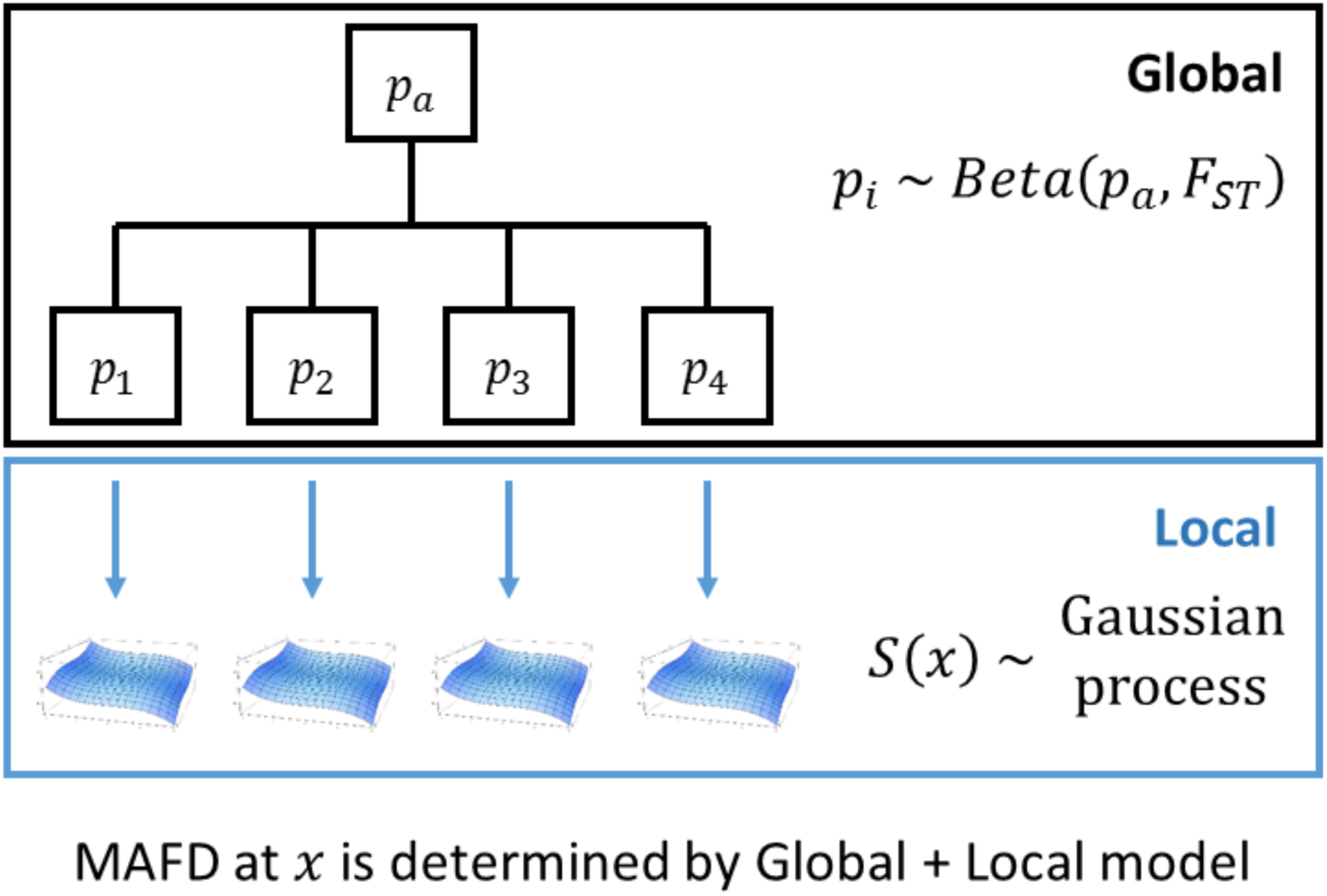
Overview of the inference levels. The Global step operates on a cluster-wide resolution, providing estimates for entire clusters based on a beta-binomial model of allele frequencies. The Local step operates within clusters, providing localized estimates across the ancestry space spanned by the individuals in each cluster. This step models allele frequencies as spatial processes operating within clusters. The Global and Local inference modules complement each other, the former picking up larger fluctuations in allele frequencies, and the latter generating a fine map that would otherwise have been hidden by the strong signal at the Global level.

**Figure 3:**
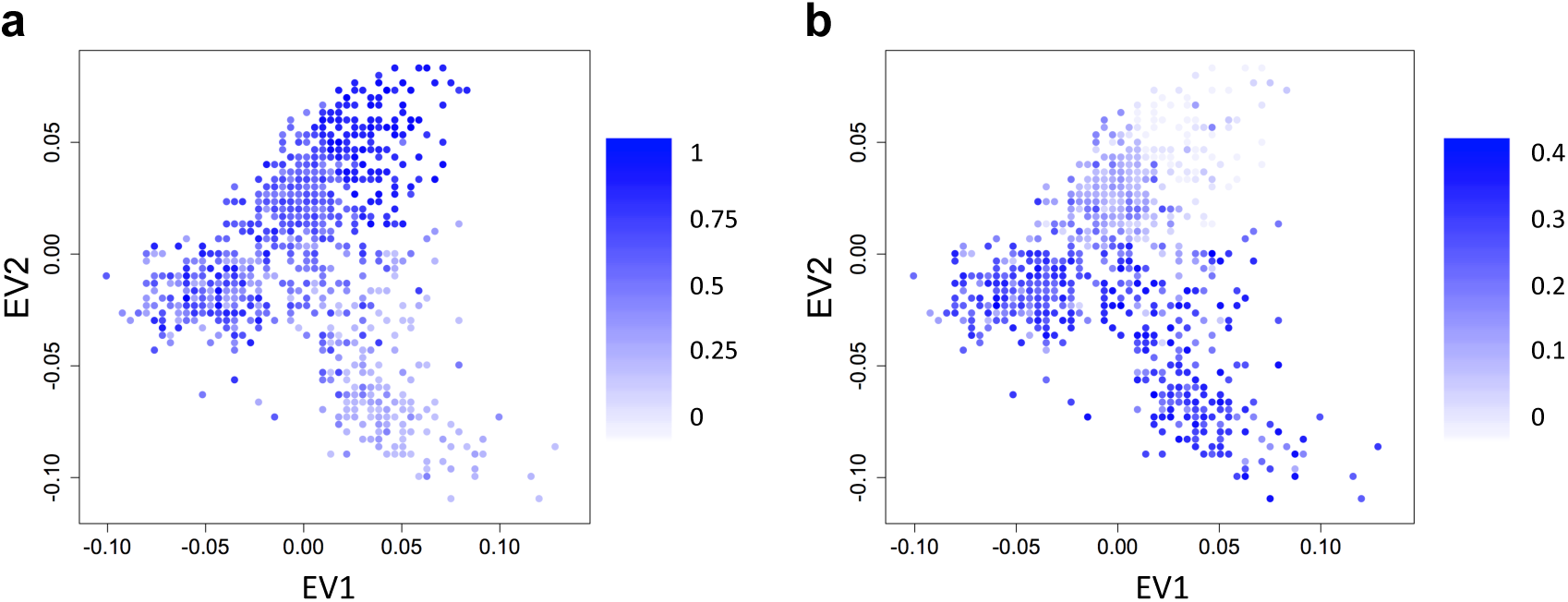
Clines detected by UNICORN in the POPRES data for two SNPs under strong selection. Intensity of color displays allele frequency estimates that vary smoothly across the map. (a) Cline of a SNP within the LCT region (lactase persistence); (b) Cline of a SNP within the OCA2 region (hair, skin and eye color).

Conceptually UNICORN aims to use as many control samples as justifiable, based on ancestry, to estimate the MAFD associated with each case sample. To motivate this model, consider two different matched case-control studies: one with equal numbers of cases and controls and the other with ten controls for each case. In the first instance, the statistical power is driven equally by cases and controls; for the latter, the number of cases is the key determinant for power. For UNICORN, the matching of controls to cases is determined by how many controls are located near each case in ancestry space. Regardless of the number of cases and controls, if there were very few controls similar in ancestry to cases, any test will have a large variance and little power. Alternatively, if there are many controls that are close in ancestry space to each case, then the variance of the test will be dominated by the case sample size. UNICORN seeks to achieve power by using information on allele frequencies from a very large sample of controls.

### 2.2 Ancestry Mapping via GemTools

Dimension reduction techniques such as principal-component analysis (PCA) are traditionally used to model complex genetic structure and to control for population stratification [11, 12, 18, 19, 17, 20]. These approaches often require many dimensions to describe the ancestry space, and this is not ideal for downstream steps of UNICORN. Instead, our algorithm first discovers clusters of subjects with relatively homogeneous ancestry, which then require fewer eigenvectors to represent ancestry within a cluster. To achieve this purpose we use GemTools [13], a software tool based on a spectral graph approach [10] quite similar to PCA. We note, however, that many popular ancestry mapping techniques could be successfully paired with UNICORN in place of GemTools.

A first step in the UNICORN algorithm involves plotting both case and control samples onto a common ancestry map without data exchanging hands (Figure 1). This is achieved by generating an ancestry map using a publicly available repository, called the “base sample”, and then projecting cases and controls onto this map via the Nyström approximation [21, 22, 23, 24]. When samples are projected onto a given ancestry map, it accurately reflects their ancestry only if the base sample spans the full range of ancestries included in the new samples [21]. Individuals with unrepresented ancestry will be projected into the available range and they will be falsely represented as more similar to the base sample. Thus, as with any genetic association study, the case collection should be restricted to samples with ancestry similar to the available control samples.

The aim of the spectral graph approach is to obtain a useful eigenmap of the genetic ancestry present in a sample. The population is represented as a weighted graph with vertices denoting individuals and weights denoting genetic similarity. Define the matrix *Y* such that *y*_*ik*_ is the minor allele count for the *i*^*th*^ subject at the *k*^*th*^ SNP. Center and scale the columns of *Y*. Instead of proceeding with computing eigenvectors and eigenvalues of *YY*^*t*^ define the weight matrix *W* as 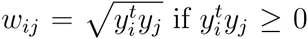 and 0 otherwise for similarity between the *i*^*th*^ and *j*^*th*^ subjects. Setting a threshold on *YY*^*t*^ to guarantee non-negative weights creates a skewed distribution of weights, so the choice of a square-root transformation leads to more symmetric distributions. This transformation also increases the robustness to outliers. Let the degree of vertex *i* be 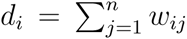 and define the diagonal matrix *D* = diag(*d*_1_,…, *d*_*n*_). The normalized graph Laplacian matrix for *W* is defined as 1 – *L*, where *L* = *D*^‒1/2^*WD*^‒1/2^. Let *v*_*i*_ and *u*_*i*_ be the eigenvalues and eigenvectors of 1 – *L* and let *λ*_*i*_ = max{0,1 – *v*_*i*_}. We can then map the *i*^*th*^ subject onto an *s*-dimensional ancestry space according to: 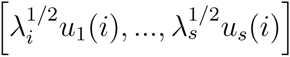. See [10] for further details.

GemTools builds on this spectral graph approach to construct eigenmaps and provide a hierarchical clustering of individuals based on ancestry [13]. To speed up computation, it is useful to avoid the cost of calculating the inner product matrix *YY*^*t*^ and then performing a spectral decomposition on a large matrix. GemTools uses a divide and conquer approach that clusters individuals of similar ancestry and then finds eigenmaps for each cluster. Homogeneous clusters of individuals are derived via Ward’s k-means algorithm. In addition to reducing computation time, this approach focuses on fine scale structure across clusters, leading to more informative maps than those resulting from a brute force computation of a single eigenmap of the entire dataset [10, 14].

New subjects are mapped onto an existing map via Nyström projection. Let *Y* represent the scaled and centered allele count vectors for the initial *n* subjects. Let *z* be the scaled allele count vector of a new individual we wish to project. We define the edge weights between the new subject and an existing individual as 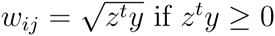 and 0 otherwise. The vertex degree of *z* is 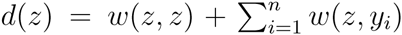. Then the eigenvector coordinates of *z* for dimensions *k* = 1,…, *s* are 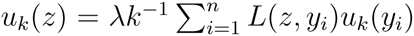 where *L*(*z*,*y*_*i*_) = [*d*(*z*)*d*(*y*_*i*_)]^‒1/2^ *w*(*z*,*y*_*i*_). Nyström projection plays a critical role in UNICORN since it allows two datasets to be mapped to the same ancestry space without the need for data sharing.

To highlight the importance of choosing a representative base sample, we estimate the eigenvectors using two different base samples derived from POPRES [15] and HGDP [25] European samples (Figure 4) [21]. When using the HGDP populations as a base (4a), the axes do not differentiate the POPRES sample. Rather the points clump together in the center of the eigenspace because their differences are dwarfed by the differences in the more diverse HGDP sample. Likewise, we found that when using the POPRES sample as a base (4b), the axes do not capture the strong differences in the highly diverse HGDP data.

**Figure 4:**
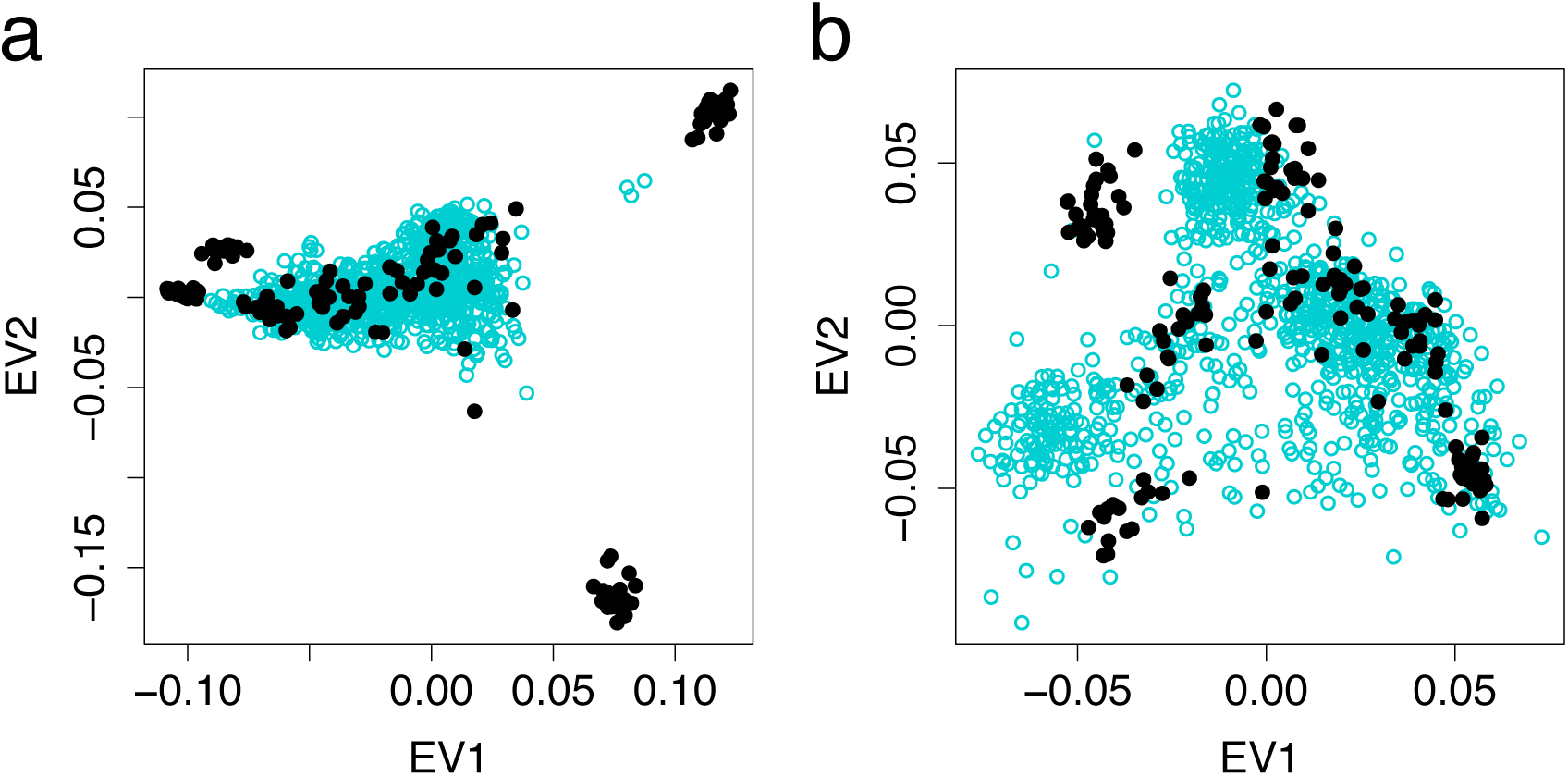
Importance of the choice of base sample for ancestry maps. When projecting new samples onto an existing ancestry map it is crucial that the base sample spans the full range of ancestries present in the new samples. If the projected samples contain unrepresented ancestries, they will still be mapped onto the ancestry range of the base set, thus distorting of their true background and leading to strongly heterogenous clusters that do not accurately reflect the allele frequencies of the new samples. (a) Base = HGDP, projected = POPRES. In this scenario we get poor resolution of ancestries in the POPRES sample. This set projects as a clump, since it looks very homogeneous relative to the more diverse HGDP base set. (b) Base = POPRES, projected = HGDP. In this scenario, the HGDP ancestries not present in the POPRES base set are still projected within the POPRES ancestry range.

### 2.3 Cluster-wide Inference

UNICORN estimates ancestry-specific allele frequencies using an efficient, flexible semi-parametric model. Frequencies are modeled in two stages to account for global and local structure. In the first stage, the data are partitioned into approximately homogeneous ancestry clusters based on eigenanalysis [14]. Next, each of these clusters is subsequently described by a secondary eigenanalysis that models local ancestry within a cluster. In stage two, local variability is modeled over the ancestry space using a Gaussian spatial process. The key to modeling local variation in allele frequency is to obtain a parsimonious representation of the ancestry not unlike a geographic map. GemTools recursively partitions the subjects until the clusters are approximately homogeneous, as judged by the leading eigenvalues [14]. Consequently two eigenvectors are sufficient to describe the residual ancestry differences within clusters at the final stage.

Each stage of the model is amenable to a simple statistical model that accounts for allele frequency variation over the ancestry space and records variability in the allele frequency estimate. In the first stage, the allele counts are modeled using a beta-binomial model with variance a function of the well-known genetic parameter *F*_*ST*_. Assume we have a data set in which GemTools detects *n* subpopulations. At this stage we want to find good estimates of the true cluster-wide allele frequencies *p*_*i*_. We model each of these frequencies as

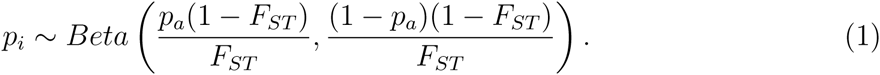

Following an empirical Bayesian setting, we use equation 1 as a prior for the cluster-wide allele frequency and use the data to guide us in selecting appropriate values for the two hyperparameters *p*_*a*_ and *F*_*ST*_. Let *p̂*_*i*_ be the average allele count in cluster *i*. Although *p̂*_*i*_ is an unbiased estimator of *p*_*i*_, it can have a large variance if few individuals reside in the cluster. From equation 1 we have

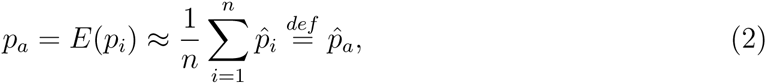

and

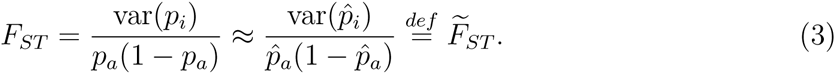

This estimator can be improved by taking into account linkage disequilibrium, the tendency of nearby alleles to descend from the same ancestral chromosome. The *F*_*ST*_ of nearby alleles must thus be similar, creating a smooth *F*_*ST*_ function across the genome. However *F̃*_*ST*_ can exhibit excessive variation which is alleviated by local smoothing through kernel regression based on genomic location.

Using equations 1-3 we estimate the prior for *p*_*i*_ through

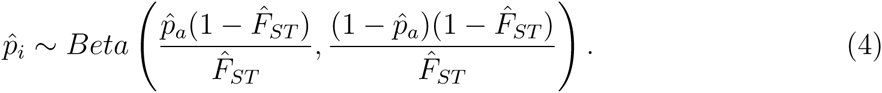

Assume we observe the genotype vector ***y*** for the *n*_*i*_ individuals located in cluster *i*. Then the posterior distribution of *p̂*_*i*_ is

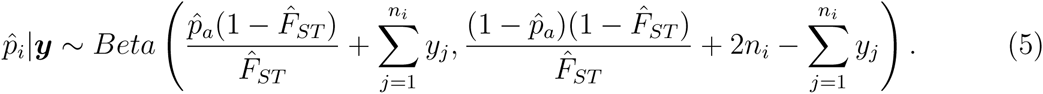

This is the distribution for the cluster-wide allele frequency that we will proceed to use for local inference.

From equation 5 it follows that the posterior mean of *p̂*_*i*_ is

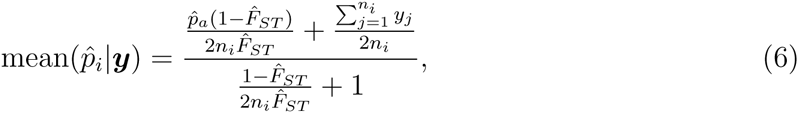

the posterior variance is

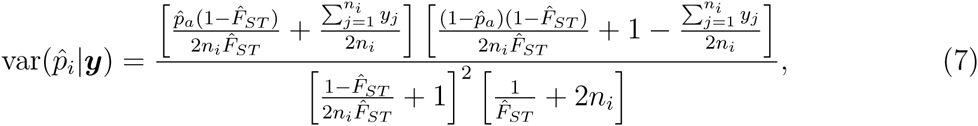

and the posterior *p̂*_*i*_|***y*** is a consistent estimator of the true minor allele frequency *p*_*i*_.

### 2.4 Within-cluster Inference

In the second stage, local structure is quantified using models made popular in the geo-statistics/kriging literature [26]. To describe the model we require the following notation: *Y*(*x*) = minor allele count at location *x* in the eigenspace; *P*(*x*) = minor allele frequency at location *x*; *S*(*x*) = deviation from cluster-wide average allele frequency at location *x* (spatial structure); *β* = cluster-wide log odds of minor allele frequency; *σ*^2^ = variance of the stationary Gaussian process (SGP); and *ϕ* = rate at which the correlation *ρ* between values of *S* at different locations decays with increasing distance *u*. Kriging methods consider a stochastic process *S* = {*S*(*x*) : *x* ∈ ℝ^*p*^}, called the signal, whose realized values are not directly observed. We do observe *Y*, the vector of allele counts, which are located in the eigenspace indexed by *x*. We assume that the distribution of *Y*(*x*) depends on *S*(*x*) and that the allele counts are a noisy version of *S* for a given set of locations *x*_*i*_, *i* ∈ 1,…, *n*. The goal is to predict S(x) at new locations where the cases have been sampled. To model local structure in an ancestry space we assume that deviations from a cluster-wide average follow a stationary Gaussian process with mean 0 and a covariance structure that will be inferred from the data. Consider the Bayesian kriging setup:

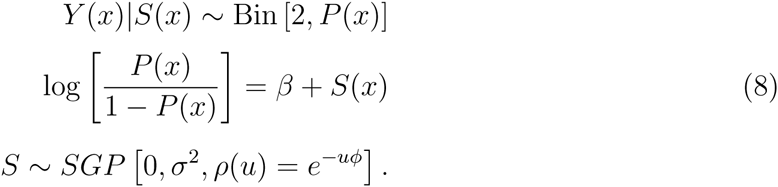

This model is appropriate because it is flexible enough to accommodate smooth allele frequency fluctuations with varying degrees of spatial correlation. With this two-stage model we can make use of our hierarchical clustering and at the same time adapt local inference to the variability present in the data, all in a Bayesian framework. Inferences are performed via Metropolis-Hastings. Ultimately, the distribution of the MAF is well approximated by a function that captures the mean and variance of the estimate.

The variance parameters of UNICORN, *σ*^2^ and *ϕ*, determine how fast allele frequencies fluctuate over the ancestry space. We can use the available ancestry space to make an informed choice of priors for these parameters. To extract the necessary variability information from the data we use a well-established method from the kriging literature: the variogram [26]. The theoretical variogram *γ*(*x*,*y*) describes the spatial dependence in a random field:

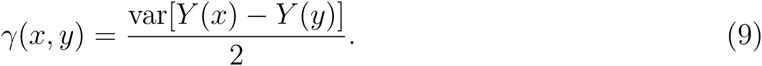

If the random field is stationary and isotropic, which is assumed here, then the theoretical variogram can be rewritten as:

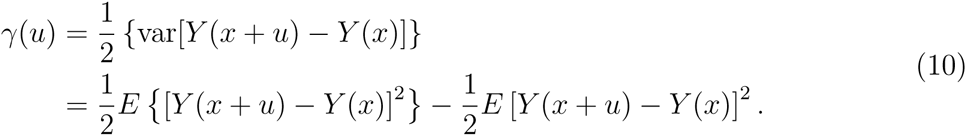

If *E*[*Y*(*x* + *u*)] = *E* [*Y*(*x*)], thus under the assumption that there exists no spatial structure, the theoretical variogram is routinely estimated via the empirical variogram:

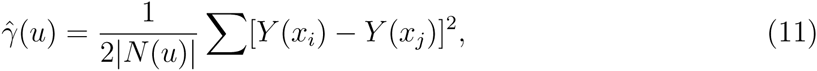

where the sum is over *N*(*u*) = {(*i*, *j*) : *x*_*i*_ – *x*_*j*_ = *u*} and |*N*(*u*)| is the number of distinct elements of *N*(*u*). But since we expect spatial structure to be present, we cannot compute the empirical variogram via equation 11 directly. Instead we estimate the spatial structure in *Y* first through linear regression in the ancestry space, and then we use the residuals and equation 11 to compute a residual empirical variogram. The next step uses both the theoretical and empirical variogram to derive values for the variance parameters. Because an algebraic expression of the theoretical variogram is complicated, we use simulation to find appropriate estimates of the variance parameters. Priors for *σ*^2^ and *ϕ* are then chosen so that their mean equals the value derived from the variogram analysis.

### 2.5 Association Test

Each case sample is mapped to a cluster in the hierarchical tree and an ancestry position x within the cluster. Combining the results from our cluster-wide inference and within-cluster inference (Figure 2) we can obtain the MAFD for this case

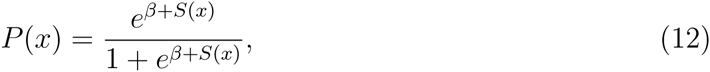

where *S*(*x*) is the spatial structure determined via the Gaussian process model (within-cluster inference) and *β* is based on the beta-binomial model (cluster-wide inference). Specifically, this expression determines the mean, *E*[*P*(*x*)], and the variance, var[*P*(*x*)], of the MAFD(*x*) which is required to perform an association test.

For an association study we sample minor allele counts [*Y*(*x*_1_),…, *Y*(*x*_*n*_)] for a sample of *n* cases. Under the null hypothesis (no association) we assume that *Y*(*x*) ~ Bin(2, *P*(*x*)). It follows that *E*[*Y*(*x*)] = *E*[*E*[*Y*(*x*)|*P*(*x*)]] = 2*E*[*P*(*x*)] and

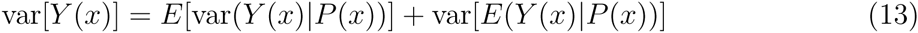

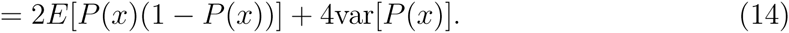

The null distribution of *Ȳ* follows from the central limit theorem:

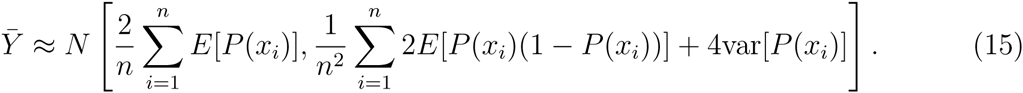

Z-scores and subsequently p-values can be computed for association tests based on equation 15. This result shows that if many controls become available for each case (decreasing var[*P*(*x*)]), then the variance of the null distribution will be dominated by the binomial sampling variance in the cases. In this setting the statistic reduces approximately to a test comparing the minor allele frequency in cases to a known population quantity and the power of the test is largely determined by the number of cases sampled. At the other extreme, if only one matched control is available for each case, then the statistic is equivalent to the usual 2-sample test and has twice the variance attainable by UNICORN with a large sample of controls. Provided the cases are well matched to a large UNICORN control sample the power can be approximated using a genetic power calculator with control:case ratio set suitably high, say ten.

## 3 Results

### 3.1 Analysis of POPRES Data

To illustrate UNICORN we use data from POPRES [15], from which we selected 160,000 high-quality SNPs (MAF > 1% and less than 1% missing genotypes) and 1000 individuals of European ancestry (each subject must have no more than 1% missing genotypes). The hierarchical ancestry structure was determined via GemTools, yielding an ancestry map that approximates the geographic map of Europe [27, 16].

For any particular study we expect the UNICORN repository will include 10-20 times as many control samples as cases. Moreover it is likely that only a fraction of these controls will be suitably matched in ancestry to the cases. Thus to mimic the realistic performance of UNICORN using the POPRES data, we needed to select a small case sample with a particular regional distribution. Specifically we randomly selected 60 POPRES samples of French and Swiss ancestry to serve as cases (6% of the total). For this constructed case-control sample, we simulated causal variants of varying allele frequencies and odds ratios. We first performed a matched-control association test, where the selected case individuals were matched to the nearest controls in the ancestry space. We then analyzed the simulated variants via UNICORN and found that it delivered more powerful results even when compared to a standard case-control association test comparing 60 cases to 600 ancestry matched controls (Figure S1).

### 3.2 Application to IBD Data

The large meta-analysis study of Crohn’s disease (CD), including 5,956 cases and 14,927 controls [28], provides a realistic test of the validity and power of the UNICORN approach. This study is perfect for detailed investigation for two reasons: first, it provides a very large sample of data that include the challenges of genotypes imputed across multiple arrays; and second, all SNPs with moderately promising signals were genotyped for 75,000 individuals in a validation study to reveal the true risk status of many SNPs.

To assess the performance of UNICORN we performed two experiments. (1) A direct comparison between UNICORN and an analysis of the full set of cases and controls. In this experiment we learn if UNICORN efficiently utilizes all the data in the control sample by comparing the power of the two tests. We do not expect UNICORN to have greater power, but we can determine if it loses power compared to a direct analysis of the data. To determine if UNICORN produces false positives we permute case and control labels and look for deviations from the expected null distribution. (2) We mimic a realistic application of UNICORN by focusing on a particular study within the larger sample, complete with ancestry matched cases and controls. In this experiment we compare performance of a direct analysis of the matched case-control study to UNICORN applied to the same cases but with the full unselected sample of controls, excluding the matched controls.

#### Experiment 1

To obtain a baseline for power in the CD dataset we performed a traditional logistic regression analysis on the full sample of cases and controls, adjusting for ancestry using principal components (LRegr). For comparison UNICORN used the full sample of controls to construct the MAFD for each case, and then performed an association test using all cases. The results for the two methods were extremely similar (Figure 5a); notably all SNPs that yielded significant results for LRegr (*p* < 5 × 10^‒8^) also yielded significant results for UNICORN. This shows that in spite of the fact that UNICORN only handled the control data indirectly via the MAFD, it maintains full power to detect associations signals. Moreover, each of the significant SNPs was also significant in the validation study [28].

**Figure 5:**
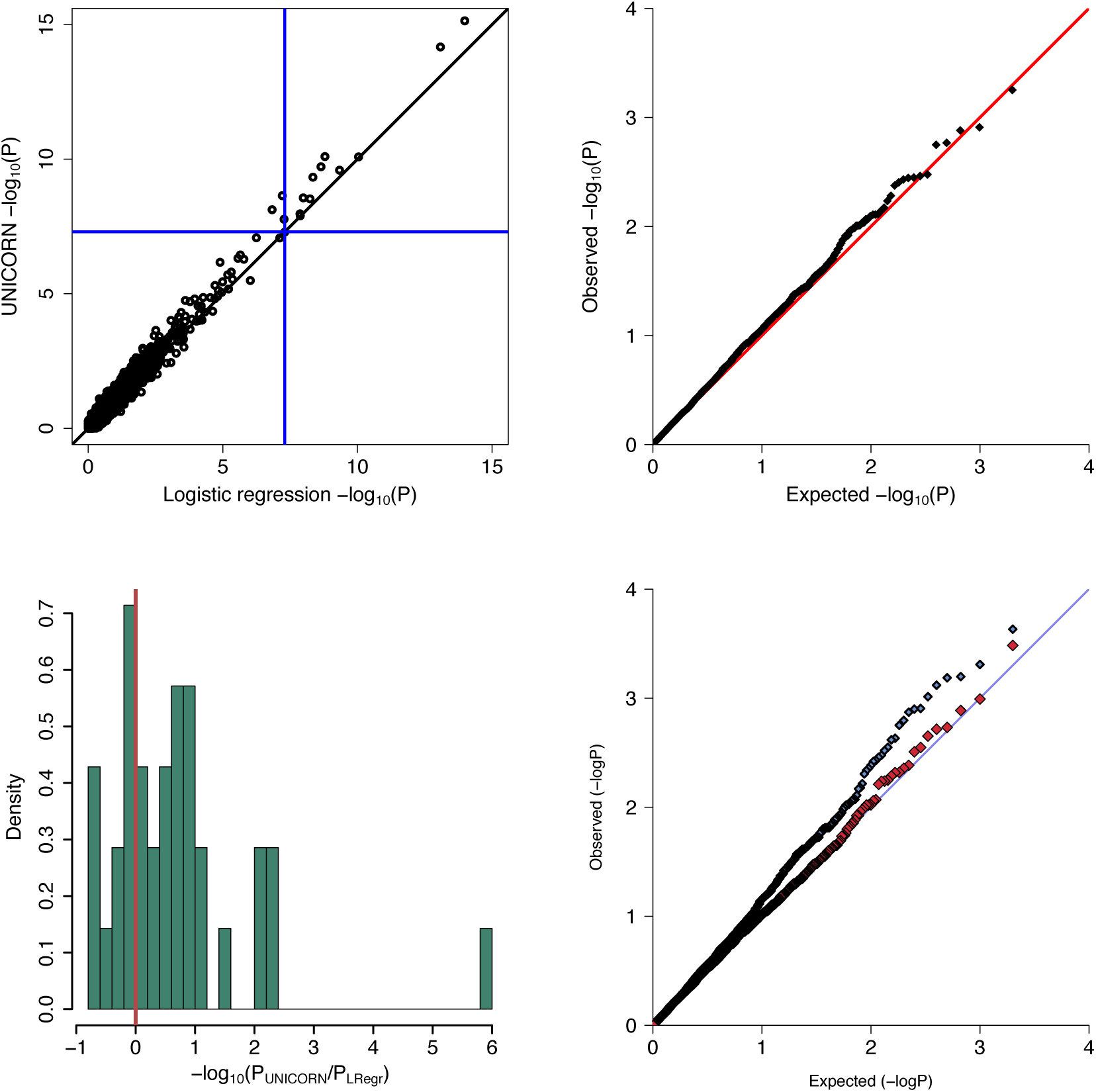
IBD analysis using UNICORN vs. logistic regression controlling for ancestry (LRegr). (a) Comparison between UNICORN and LRegr on the full 7-study CD dataset. All significant SNPs detected by LRegr were also significant in UNICORN, and each of these SNPs was significant in the validation study as well. (b) UNICORN null distribution obtained by permuting affection status in the full case-control dataset. The resulting distribution of p-values produced by UNICORN is well calibrated, indicating a good control of false positives. (c)-(d) UNICORN applied only to cases from the Belgian study using all controls excluding that study. (c) Difference in p-value magnitude between UNICORN and LRegr applied only to Belgian case-controls. Results are shown only for SNPs that were found significant in the validation study [28]. All SNPs showing a substantial differences favored UNICORN, particularly the SNP that had the highest signal in [28]. (d) P-P plot for UNICORN (blue) compared to the null distribution with permuted phenotype labels (red). The blue P-P plot shows some signal was detected and the red P-P plot shows that UNICORN yields an appropriate null distribution when there is no signal present.

To examine the overall validity of the tests we computed the λ_1000_ genomic control factors [29, 30] and found both tests performed well: λ = 1.03 for UNICORN and λ = 1.02 for LRegr. To further evaluate the validity of UNICORN in the absence of polygenic effects we permuted the case and control labels to remove association [31]. The distribution of p-values produced by UNICORN is well calibrated to meet null expectations (Figure 5b) and the genomic control factor for this distribution is λ = 1.01. In total this experiment shows that UNICORN makes efficient use of the full data without inducing false positives.

Finally, to illustrate the impact of each level of population structure we analyzed these data 3 ways: (1) ignoring the effect of ancestry altogether; (2) modeling only the global structure using the first level of UNICORN; and (3) modeling the global and local structure with UNICORN. As expected, not account for ancestry leads to a P-P plot with strong evidence of overdispersion; incorporating the global level of UNICORN leads to a marked improvement in the distribution of test statistics; and finally modeling additional structure at the local level leads to even greater reduction of false positives (Figure S2).

#### Experiment 2

UNICORN is designed to permit analysis of a case-only sample by utilizing controls drawn from a repository. To evaluate performance in this setting we extracted the IBD-CD Belgian study for further investigation. This study consists of a sample of 666 CD cases and 978 controls of similar ancestry. A case-only sample applying UNICORN would have access to all 14,927 controls minus the 978 Belgium controls. For comparison we contrasted the results of analysis of this well matched study using LRegr with UNICORN using all non-Belgium controls.

Not surprisingly no SNP is genome-wide significant for this modest sample of cases for either analysis. To compare the power, we evaluated the behavior of the association tests for the 163 SNPS that showed genome-wide significance in the validation study [28]. Taking this as truth, we favor whichever method yields a smaller p-value in the comparison (Figure 5c). For 70% of these loci the evidence for association from the UNICORN analysis was stronger than the LRegr analysis, and for the 30% where it was not superior the tests were nearly identical for both approaches with neither test showing a signal. One surprising result was that the most highly significant SNP in the validation study exhibited a p-value 6 orders of magnitude smaller with UNICORN than LRegr (Figure 5c). This SNP has a relatively large *F*_*ST*_.

Based on the P-P plot, UNICORN p-values detect a modest signal for many SNPs. To assess the validity of the test we permuted the case and control labels to remove association and found that the overall distribution of the UNICORN test was appropriate (Figure 5d, red). This experiment supports the great potential of UNICORN to increase power without incurring false positive findings.

### 3.3 Detection and Removal of False Positives

One of the major challenges in the analysis of genetic data is controlling for the technical variability across different SNP arrays, imputation pipelines and genotyping approaches. This challenge is equally great when applying UNICORN; however careful attention to process and quality control (QC) can greatly enhance the reliability of the analysis.

Ultimately the UNICORN repository will consist of an assimilation of samples from tens, if not hundreds, of individual studies. Hence it will certainly include multiple SNP arrays. To avoid exacerbating study-specific biases, all samples in the repository will be imputed using a common pipeline. As proof of concept, imputation was performed jointly for the CD controls used here, which stem from seven different studies and arrays. After first performing the QC procedures described below, no significant array bias was detected in the study [28]. The IBD study demonstrates that a homogeneous control collection can be assembled from different sources, provided care is taken with the imputation and QC. Likewise, imputation on the cases should follow the same pipeline as implemented for UNICORN controls.

Based on our investigation of challenges due to imputation and array-based biases we have identified a reliable approach that is quite similar to the technical QC and assay evaluation in use for routine genetic analysis. The objective is to identify SNPs with unusually small p-values relative to their linkage disequilibrium (LD) neighbors. Such signals are almost always due to technical artifacts. SNPs with p-values not supported by their LD neighbors can be identified using either nonparametric regression or a hidden Markov model via DIST [32]. Both procedures successfully flag SNPs with outlier p-values.

To illustrate this approach we used individuals with European ancestry from the HGDP dataset as cases in comparison with the CD controls as part of a UNICORN association analysis. Any signal detected by such a test stems from technical artifacts and should be flagged as such. Prior to QC, UNICORN did indeed return some signals on chromosome 1 (Figure 6a,b). We ran a nonparametric kernel regression with binwidth of 2Mbp across the series of –*log*_10_ p-values and flagged results as noise if the smoothed value differed from the actual value by more than one order of magnitude. This procedure eliminated all the isolated signals (Figure 6a,c).

**Figure 6:**
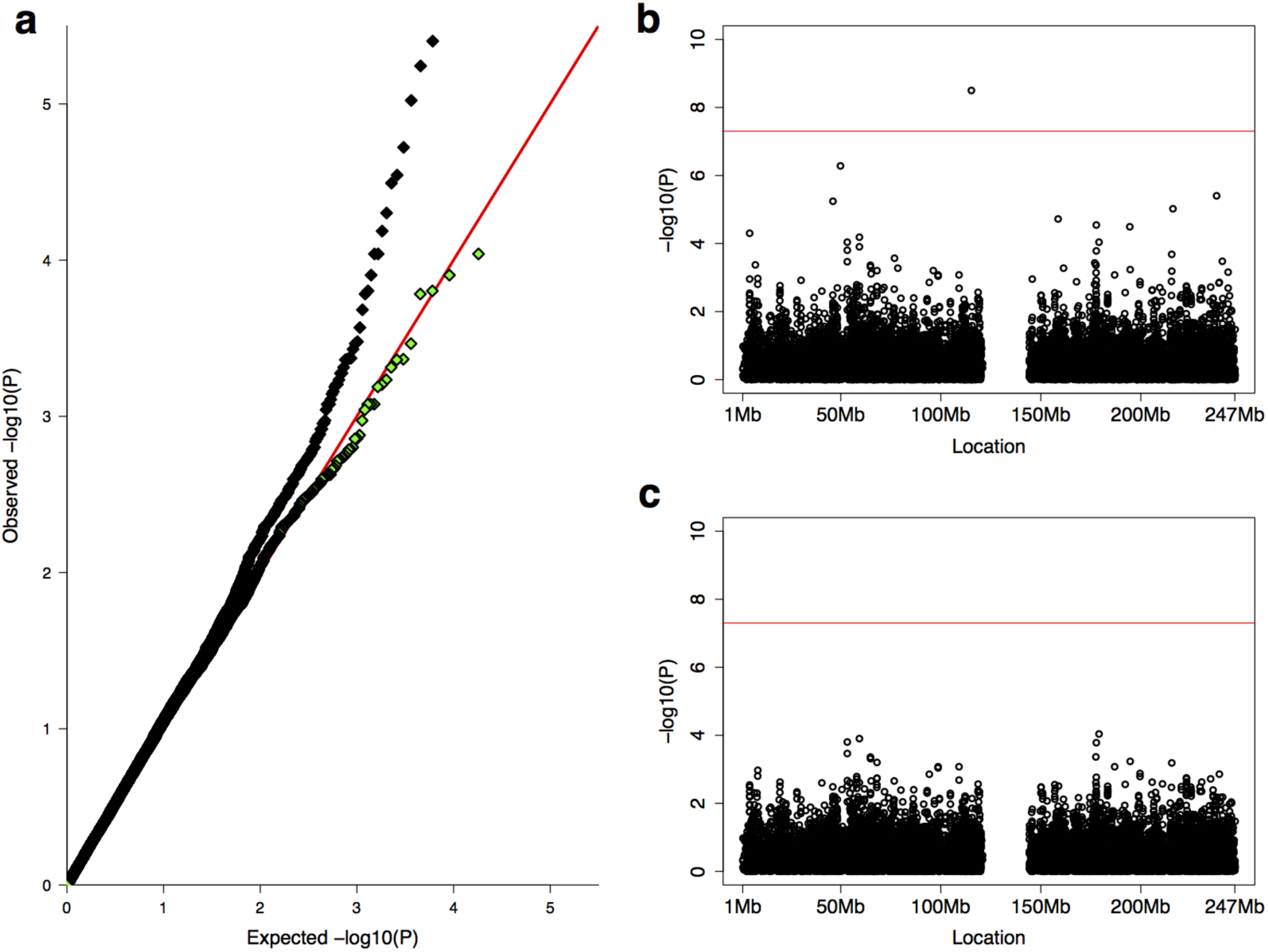
Detection and removal of false positives using nonparametric smoothing. We created a UNICORN study by using individuals selected for European ancestry from the HGDP dataset and comparing them to the CD controls. Any signals in this comparison are likely due to technical artifacts. (a) P-P plot of UNICORN results before (black) and after smoothing (green) to reduce noise. Notice the strong presence of signal in the black P-P plot despite the expectation of no signal when comparing two control data sets. (b) Manhattan plot of UNICORN p-values before smoothing exhibits isolated signals without support in the immediate LD neighborhood. (c) Isolated signals are removed after smoothing.

Another precaution can be employed to remove false positives due to differing SNP arrays. A comparison between UNICORN controls and controls measured on the same array as the cases should reveal SNPs that cannot be reliably compared across these arrays. Any SNPs exhibiting a signal in these experiments should be removed from further investigation. Moreover, SNPs identified by internal comparisons across chips in development of the UNICORN repository will be noted on the UNICORN web site.

In conclusion, we note that similar to any large genetic association study, UNICORN can yield false positives due to technical artifacts. This challenge arises in part because UNICORN requires imputation in the control data sets to obtain a common set of SNPs across arrays for subsequent analyses. When the case sample is genotyped on an array that is not well represented in the control sample the challenge is greater; however, we have found that post analysis cleaning can remove false positives that arise.

## 4 Discussion

An essential feature of a genetic association study is a large control sample, chosen to represent the case sample in ancestry [33, 34, 35]. Although suitable control samples can sometimes be obtained from public repositories, it typically requires substantial analytical effort to process controls along with the case sample. The goal of UNICORN is to obviate this need, at least partially, by automatically providing equivalent information from controls collected previously, for example controls deposited in dbGaP. For each case sample, the algorithm uses available controls to estimate the allele frequencies matched by ancestry. In this way UNICORN will facilitates case-control studies, even if the study has characterized only case samples, and thereby optimize the discovery of risk variants. By providing a control sample that is ancestrally matched to cases, without requiring resources or effort from the user, UNICORN provides advantages even for case-control studies for which a set of control samples has already been collected. In our proposed implementation, the end user would not experience significant compute time because the MAFD can be pre-computed and easily queried based on the user’s cases.

Not sampling controls as part of the study design precludes the direct inclusion of covariates in the analysis. In some settings a properly chosen covariate can greatly enhance power, while in other scenarios covariates can reduce power [36] or bias the analysis [37]. Even in the former setting when covariates are useful, UNICORN can provide a more powerful analysis due to the enhanced estimate of the population allele frequency derived from a much larger sample of controls (Figure S3). Moreover, if covariates are measured in the cases, it is possible to perform conditional genetic analysis on subsets of the data using UNICORN. Such an analyses contrasts the SNP allele frequencies in a subset of the cases with the estimated population allele frequency of ancestry-matched controls.

A population control sample by definition includes some subjects that should be classified as cases. This will reduce power, but it will not generate an excess of false positives, and the impact on power increases with the frequency of the disorder under investigation. When screened controls have not been collected, however, it is common practice to rely on population controls and UNICORN has no special weaknesses or strengths with regard to this issue.

The current analysis features samples of European ancestry, but the framework is applicable to other ancestries as well. Due to its multiple levels of inference, UNICORN can accommodate populations of quite complex structure, such as that found from African populations [38], as well as the simpler structure of European populations. Our experiments suggest that UNICORN models ancestry as effectively as PCA, hence, we expect it will perform well in other ancestries. Analyses of recently admixed populations are more challenging, however, and will require new additions to the UNICORN methodology.

In addition to the potential gain in power, UNICORN also has the potential to strengthen subject privacy. The ability to identify an individual from their anonymous genetic information in a public database threatens the principle of subject confidentiality [39]. Knowledge of an individual’s genotype at relatively few SNPs is sufficient to uniquely identify a person; indeed this is the basis of DNA forensics. Protected repositories such as dbGap exist so that genome-wide data may be shared among responsible parties without exposing subjects to a loss of privacy. But a second level of privacy loss is also of concern. Based on reported allele frequencies in cases and controls, given a very large number of SNPs, it is possible to determine with high probability if an individual is a case or control in the study, or not in the study at all [40, 41, 42]. By restricting the exchange of genetic data to ancestry coordinates, UNICORN could overcome both of these challenges. Additionally, our grid-based approach, where we return frequency estimates from the pre-computed grid point closest to the case instead of the actual case location, provides another layer of security for the control identities by adding a small degree of randomness to our predictions.

Results from the Exome Aggregation Consortium (ExAC) motivate this work. ExAC has made substantial progress toward the goal of assembling exome sequencing data from a variety of large-scale sequencing projects to make summary data widely available, with over 60,000 individuals worth of data available [43]. Currently, ExAC provides allele frequency information for these samples. Through UNICORN, we aim to enhance this concept by generating ancestry matched MAFD estimates for additional subjects. Although there are more technical challenges involved with sequence data than genotyping arrays, the ExAC project provides support to the belief that these data can be successfully aggregated and harmonized for use in UNICORN. We are thus currently in the process of extending the UNICORN framework, which will require further development and refinement for rare variation.

The UNICORN database and web server are in preparation and a limited version focusing on populations of European descent is slated for release late in 2016. If successful, UNICORN will dramatically improve access to the control resources stored in repositories such as dbGaP and can also make use of control samples from the same study as well as from other studies. We predict that UNICORN will hasten the discovery of genetic variation conferring risk for disease in three ways: by providing ancestrally-matched allele frequencies; by its careful integration of data sets; and by making genetic association analysis simpler.

## 5 Appendix

The covariance between two points of the Gaussian process at distance *u* is 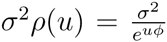. Notice that *ϕ* is the characteristic length-scale of our process: it determines how far apart two individuals must be for the allele frequency to change significantly.

Inference in this model is performed via MCMC. Write ***S*** = [*S*(*x*_1_),…,*S*(*x*_*n*_)] for the vector of values of *S* at the observed locations *x*_*i*_ and 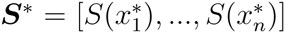 for the vector of values of *S* at the target locations 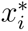 for which predictions are requested. Define ***P*** and **P*\** similarly and let ***Y*** be the genotype data at the observed locations.

A cycle of the MCMC algorithm involves first sampling from (*σ*^2^, *ϕ*)|(***Y***, ***S***, *β*), then from *S*_*i*_|(***S***_‒*i*_, ***Y***, *σ*^2^, *ϕ*, *β*) and finally from *β*|(***Y***, ***S***, *σ*^2^, *ϕ*). Here ***S***_‒*i*_ denotes the vector ***S*** without its *i*^*th*^ element. Note that since conditionally on *S* the random variables *Y*_*i*_ are mutually independent we have:

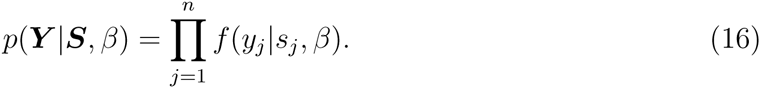

Using equation 16 we have:

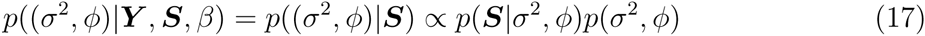

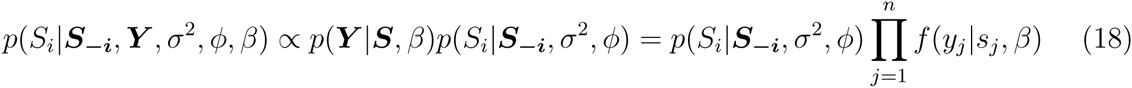

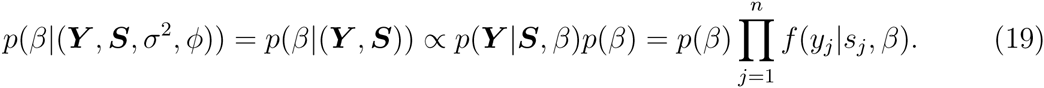

From the Gaussian process assumption it follows that *p*(***S***|*σ*^2^,*ϕ*) has a multivariate normal density (mean 0 and covariance matrix *σ*^2^*e*^−*Uϕ*^ where *U* is the Euclidean distance matrix for the locations referred to by ***S***) and *p*(*S*_*i*_|***S***_‒*i*_, *σ*^2^, *ϕ*) has a univariate normal distribution. Also, we know that *f*(*y*_*j*_|*s*_*j*_, *β*) follows a binomial distribution (with success probability 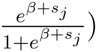 and *p*(*β*) and *p*(*σ*^2^, *ϕ*) are the priors. Being able to draw from all these distributions enables us to apply the following component-wise Metropolis-Hastings algorithm.

1. Set initial values of *β*, *σ*^2^ and *ϕ* by drawing from their respective priors. Set the starting value for each *S*_*i*_ to 0.
2. Update (*σ*^2^, *ϕ*)

- choose a new value (*σ*^2′^, *ϕ*′) from some appropriate proposal distribution *q*((*σ*^2′^, *ϕ*′)|(*σ*^2^, *ϕ*))
- using equation 17 accept (*σ*^2′^, *ϕ*′) with probability

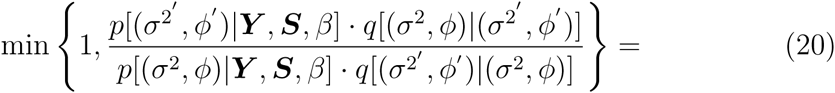

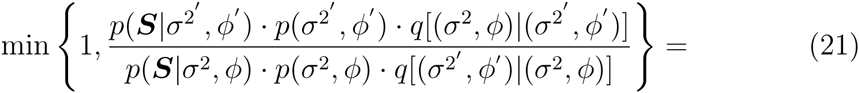

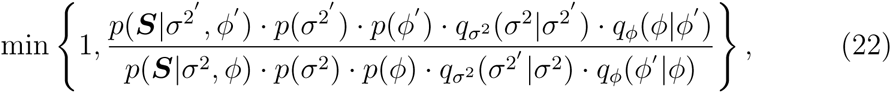

where the last equality holds if *σ*^2^ and *ϕ* are independent.
- the prior distributions for *σ*^2^ and *ϕ* as well as the jumping distributions *q*_σ^2^_ and *q*_*ϕ*_ can be gammas.
3. Update ***S***

- choose a new value Si for the 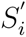 for the *i*^*th*^ component of ***S*** from the transition probability function 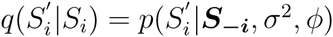
- using equation 18 accept 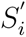 with probability

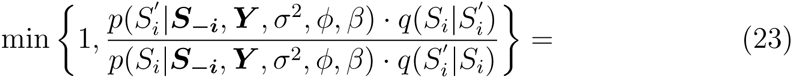

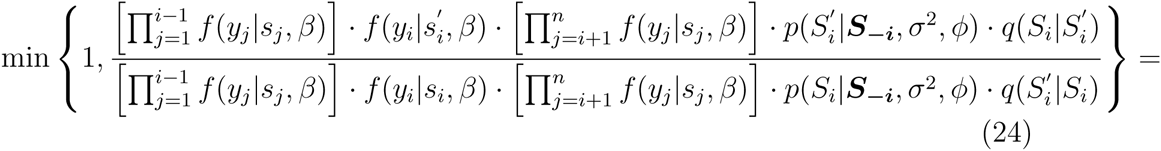

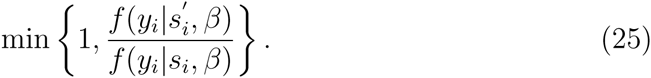
- repeat the previous two steps for all *i* = 1, …,*n* to complete updating ***S***
4. Update *β*

- choose a new value *β*′ from some appropriate proposal distribution *q*(*β′*|*β*)
- using equation 19 accept *β*′ with probability

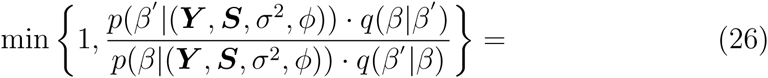

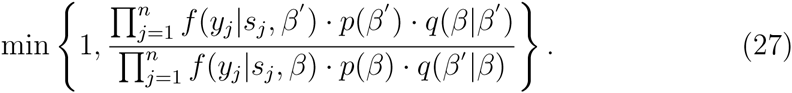
- the prior distribution of *β* is determined by the cluster-wide allele frequency inference step, and the jumping distribution *q* can be a normal distribution

Repeat steps 2-4 (with an optional burn-in period and thinning) to obtain draws from the equilibrium distributions. We are now able to draw from the posteriors of *σ*^2^, *ϕ*,*β*, ***S***. We proceed with:

5. Draw a sample from the multivariate Gaussian distribution of ***S***\*|(***S***, ***Y***, *σ*^2^, *ϕ*, *β*) where the values of ***S***, *σ*^2^,*ϕ*, *β* are those generated in steps 2-4. Using the conditional independence structure of our model this step reduces to drawing from ***S***\*|(***S***, *σ*^2^, *ϕ*). The Gaussian process assumption implies that:

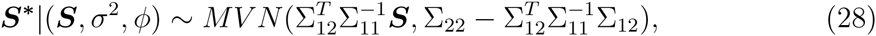

where

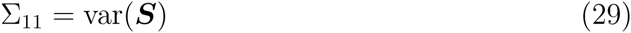

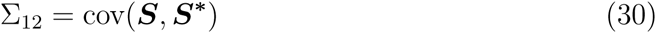

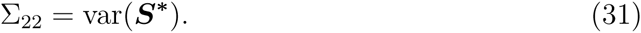 Each of these matrices can be computed based on the variance properties defined by *σ*^2^ and *ϕ*.
6. Compute ***P***\* based on the current values of ***S***\* and *β*:

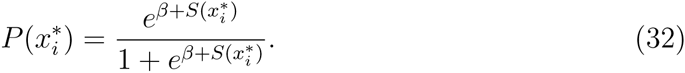

Iterating steps 5-6 gives us the predictive distribution 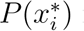 for all points at which we want to infer allele frequencies.

## 6 Supplemental Data

Supplemental Data include one figure and one list of consortia authors.

## 7 Acknowledgments

This work was supported by the National Institute of Mental Health grant R37MH057881 (B.D. and K.R.) and National Institute of Health grant MH101244-02 (B.M.N.). The authors declare that there are no conflicts of interest. We thank the referees for insightful comments.

## 8 Web Resources

The R package GemTools together with its documentation can be dowloaded from: http://wpicr.wpic.pitt.edu/WPICCompgen/GemTools/GemTools.htm

